# The Molecular Architecture of Somatic Spines of the Lateral Septum

**DOI:** 10.1101/2025.05.19.654857

**Authors:** Daniela Hacker, Erich Weisheim, Michaela Schweizer, Undine Schneeweiß, Michael Brecht, Marina Mikhaylova

**Author notes:** Corresponding author email address: Marina Mikhaylova.

## Abstract

The lateral septum is a key subcortical structure and has been implicated in kinship memory. Across species, the ability to recognise relatives is conserved and is reflected in stable, lifelong memories. The molecular mechanisms underlying kinship memory remain elusive. Here we investigate the synaptic architecture of septal circuits and focus on somatic spines, an apparent synaptic specialisation of this mainly GABAergic structure. We uncover the molecular organisation of septal somatic spines found in the lateral septum using a combination of unbiased imaging methods and confocal microscopy. We used classical label-free methods such as transmission electron microscopy and Golgi stainings and established new culturing methods for dissociated and organotypic septal slice cultures that were kept in culture over multiple weeks. We describe the presence, morphology, ultrastructure and molecular composition of excitatory somatic spines across multiple developmental stages in various model systems. While smaller than dendritic spines, somatic spines exhibited distinct features, frequently containing secretory organelles such as phagophores and endosomes, but often lack a spine apparatus and ribosomes. Our findings offer insights on the molecular architecture of septal somatic spines and establish a basis for further investigations on the somatic spines of the lateral septum and their role in kinship memory.

**Significance statement:** Spine synapses represent stable synaptic connections that allow neurons to chemically communicate with each other. Although known for decades, the unusual spine synapses found directly on neuronal cell bodies in the lateral septum have not been characterised in detail. Using traditional, unbiased approaches such as electron microscopy and Golgi stainings, we replicate the initial findings. Additionally, using fluorescent microscopy of dissociated and organotypic septal cultures, we characterise the development and composition of septal somatic spines. We find that they are excitatory, contain membranous organelles and develop independently of extra-septal input. We thus describe model systems suitable for the investigation of somatospiny neurons of the lateral septum.

## 1. Introduction

The brain is made up of various neuronal cell types and many different synapse types. Differences in their properties include the type of signal transmission, synaptic vesicle count, synapse strength, size, shape, localisation and molecular composition. The introduction of cell type-specific Fluorescence Activated Synaptosome Sorting (FASS) allowed the cell type specific analysis of synapse types (Biesemann et al. 2014; Kaulich et al. 2024; van Oostrum et al. 2023). While this technique allows for a high-throughput analysis of synapses of a specific cell type, it does not grant information on their subcellular localisation. Synaptic spine morphology plays a crucial part in plasticity and can be broadly categorised into mushroom, stubby, thin and filopodia-like ranging from mature to immature spines (Pchitskaya and Bezprozvanny 2020).

An especially rare subtype is the somatic spine synapse that do not make up the majority of any neuron’s synapses. In the lateral septum, a large proportion of spiny GABAergic neurons can be found (Robert L. Jakab and Leranth 1990b), first described by (Onteniente et al. 1987). These somatospiny neurons were suggested to be GABAergic (Robert L. Jakab and Leranth 1990a; Onteniente et al. 1987). However, the lateral septum is comprised of a large percentage of GABAergic neurons (75% to 100% of neurons) (Risold and Swanson 1997; Rizzi-Wise and Wang 2021; Wong et al. 2016; Zhao, Eisinger, and Gammie 2013), and thus does not distinguish the cell type. The septum contains a large proportion of spiny GABAergic neurons (Robert L. Jakab and Leranth 1990b) first described by (Onteniente et al. 1987). Although observations regarding the identity of these somatospiny cells and the cells innervating them has been made (Robert L. Jakab and Leranth 1990a; 1990b; R L Jakab and Leranth 1991; R. L. Jakab, Naftolin, and Leranth 1991), no systematic description exists. The somatic spines have further been suggested to receive excitatory hippocampal, vasopressinergic and catecholaminergic input (Robert L. Jakab and Leranth 1990a; 1990b; R L Jakab and Leranth 1991; R. L. Jakab, Naftolin, and Leranth 1991; Robert L Jakab and Leranth 1993).

The exact region of the lateral septum in which somatospiny neurons are found, harbours a population of neurons mediating kinship memory (Clemens, Wang, and Brecht 2020). Here, the connectivity between the septum and hippocampus mediates whether or not rats prefer the presence of their relatives (Clemens, Wang, and Brecht 2020). While younger animals prefer the company of littermates, older animals prefer the presence of non-littermates. Lesioning the lateral septum abolishes preferences (Clemens and Brecht 2021). Kinship memories are usually especially stable and in humans are even relatively resistant to aging and disorders impairing memory retrieval. This stable connection is likely encoded by a stable structure.

Generally, spine synapses are considered more stable and mature than i.e. dendritic shaft synapses (Grutzendler, Kasthuri, and Gan 2002). A structure often found in stable and mature spine synapses is the spine apparatus (Yap et al. 2020), a specialised endoplasmic reticulum-derived organelle (Spacek and Harris 1997). Spines can also contain other organelles such as ribosomes and membranous organelles which can provide the synapse with membrane and proteins to be used e.g. during plasticity-induced remodelling (Park et al. 2006). Although abundantly present in the dendritic shaft (Spacek and Harris 1997), endosomal organelles were seen in less than 20% of spines in the hippocampus (Cooney et al. 2002) despite them being required for the enlargement of spines (Park et al. 2006). GABAergic inhibitory inputs can be found along dendrites as well as on the soma and the axon initial segment (Lipkin and Bender 2023; Vereczki et al. 2016; Kwon et al. 2019).

Despite the high density of somatospiny neurons in the lateral septum, surprisingly little is known about their role. Both the developmental emergence and molecular composition of septal somatic spines remains undiscovered. Here we introduce novel methods to study somatic spines and aim to understand the development and molecular composition of a population of somatospiny neurons in a mostly GABAergic environment.

## 2. Material and Methods

### Dissociated septal neuron cultures from P0 rats

P0 Wistar (RjHan:WI) rats were decapitated and the brain and the septi were isolated from 1 mm thick coronal slices obtained from a tissue chopper. All tissue isolation steps were done in ice-cold HBSS. Once septi from all animals were collected in 10-15 ml HBSS, they were carefully washed 5x with 9 ml ice-cold HBSS under a flow-hood. For each septum, 100 µl 0.25% Trypsin/EDTA solution was added to the septi which were suspended in 2 ml HBSS. After 15 min at 37°C, Trypsin was removed by washing 5x with 9 ml 37°C warm HBSS. 2 ml remaining HBSS-tissue mixture were moved to a 2 ml reaction tube and triturated with 20G/26G needles. The cell solution was filtered (Easy Strainer, Greiner 542000) and the filter washed with HBSS. Cells were then manually counted in a Neubauer chamber and plated in densities between 40000- 80000 cells/ml in prewarmed full medium (Dulbecco’s modified Eagle’s medium (DMEM), 10% fetal bovine serum (FBS), 1x penicillin/streptomycin, 2 mM glutamine). After 1 h at 37°C and 5% CO2 the medium was exchanged to 37°C warm Brain Phys medium (Gibco) with added 1x SM1 and 0.5 mM glutamine supplements. To counteract a change in osmolarity due to evaporation of the medium, after one week 10% medium was removed and 20% fresh medium added.

### Organotypic slices culture

Organotypic slice cultures were prepared similar to (Gee et al. 2017). P5 Wistar (Ri- Han:WI) rats were decapitated and the brain isolated and placed onto sterile filter paper and slightly rinsed with dissection medium (248 mM sucrose, 26 mM NaHCO_3_, 10 mM D-glucose, 4 mM KCl, 5 mM MCl_2_, 1 mM CaCl2, 0.001% Phenol red, add Milli-Q water to 500 ml, osmolality should be 310-320 mOsm kg and pH *>* 8 (red-magenta); before use, add 1 ml kynurenic acid to 50 ml and bubble solution with O_2_ until orange). Afterwards, the brain was sliced into either 400 or 600 µm thick coronal slices using a McIlwain TC752 tissue chopper in anterior-posterior direction and carefully transferred into ice-cold dissection medium. Slices containing the septum were isolated and the septum was isolated from the slices with two cuts using tweezers. Septal slices were then transferred under a flow-hood and gently placed onto membranes that were sitting in slice culture medium (394 ml MEM, 100 ml heat-inactivated horse serum, 1 mM L-glutamine, 0.01 mg ml insulin, 14.5 mM NaCl, 2 mM MgSO_4_, 1.44 mM CaCl_2_, 0.00125% ascorbic acid, 13 mM D-glucose) at 37°C and 5% CO_2_. After placing, slices were moved to incubators at 37°C and 5% CO_2_ before new slices were isolated. Cultures were fed twice a week every 2-3 days by removing the old slice culture medium and adding 37°C warm new medium.

### Field electroporation

Organotypic slices were electroporated according to (Kawabata et al. 2004). The slice was positioned on top of the electrode on its membrane. The cathode was filled up with slice EP buffer. After positioning the slice and the membrane, a drop of 0.5 µg/µl DNA diluted in slice EP buffer was pipetted on top. The anode was placed on the liquid’s surface and a poring pulse initiated, followed by a transfer pulse as specified in the manufacturer’s protocols. The poring pulse was set to 50 V with a pulse length of 5 ms and a decay rate of 10%. The transfer pulse was set to 20 V with a pulse length of 50 ms and a decay rate of 40%. Electroporation was performed using AC.

### Microscopy - preparation

#### Immunocytochemistry (ICC) of dissociated cultures

If not described otherwise, neurons at various DIV were fixed by replacing the culture medium with 4% PFA/4% sucrose for 10 min. The fixative was then washed off 3x for 10 min with PBS. To permeabilise the cells, they were incubated with PBS containing 0.25% Triton X-100 which was removed with two five min wash steps with PBS. Coverslips were blocked with blocking buffer (PBS containing 10% heat inactivated horse serum (30 min at 65°C) and 0.1% Triton X-100). Coverslips were then incubated over night at 4°C with primary antibody diluted in blocking buffer (BB-HE). If used, the cells were incubated with Phalloidin at the same time. On the next day, the primary antibody was washed off with three 10 min PBS washing steps. Suitable secondary antibodies were diluted in blocking buffer and incubated for 1 h at room temperature. Secondary antibody was washed off three times for 10 min with PBS. When necessary, prelabelled primary antibodies were diluted in blocking buffer and added to the cells at this stage and incubated at 4°C over night. The antibody was then washed off with three 10 min PBS wash steps. Coverslips were then mounted on glass coverslips with the cells facing a mowiol droplet on the glass coverslip. Samples could be imaged ∼ three hours after airdrying them in the dark.

#### Immunohistochemistry (IHC) of cryosections

Cryosections obtained from perfused rat/mouse brains were cut with a cryostat to 30- 40 µm thickness. For this procedure, the perfused brains were incubated with 30% sucrose in PBS to prevent ice-crystal formation and then frozen at -80°C and cut at – 20°C coronally with a cryostat. Blade and stage temperature were adjusted around - 20°C max temp. -15°C min temp -25°C. Sections were incubated with primary antibody diluted in blocking buffer (PBS containing 10% heat inactivated horse serum (30 min at 65°C) and 0.1% Triton X-100) on a horizontal shaker at 4°C for at least 36 h. Sections were washed with PBS twice for a total of 1 h, followed by two to three wash steps with PBS containing 0.2% BSA for 1 h. Secondary antibody was diluted in blocking buffer and incubated with the sections for 2 h at room temperature. Sections were then washed three times with PBS. If desired, DAPI was added 1:500 in the second last wash step. For mounting, slices were rinsed in tap water and gently placed on glass slides that were airdried. A drop of mowiol was placed on the slice and a coverslip fixed on the sample.

#### IHC of organotypic slices

PFA-fixed organotypic slices as in 3.1 Fixation were permeabilised with 0.4% tritonX- 100 in PBS for 30-90 min. After three washes with PBS, the slices were quenched using 50 mM NH_4_Cl followed by three washes with PBS and blocking for 1 h. Primary antibody diluted in blocking buffer and slices were incubated for 72 h at 4°C. After three washes with PBS, secondary antibody was diluted in blocking buffer and incubated with the slices overnight. Slices were then washed three times with PBS and optionally incubated with DAPI 1:500, before another PBS wash step. Slices were mounted using mowiol.

#### Golgi staining

Brains were stained according to FD Rapid GolgiStain Kit - FD NeuroTechnologies. In brief, P14 brains were incubated in impregnation solution for circa two weeks. During this incubation time, the impregnation solution was exchanged once. Afterwards, the brains were washed twice with protection solution for up to one week. To prepare the brains for freezing, they were incubated in cryoprotection solution (30% sucrose in PBS). They were then placed at -80°C and stored until they were sectioned with a cryostat. They were mounted on gelatin-coated slides using protection solution. Slides were thoroughly washed with tap water for 1 h and additionally for 10 min with dH_2_O. They were coated by incubating for 1 min with a 1% gelatine, 0.1% chromalaun solution. They were stored vertically at room temperature protected from dust until further usage. After airdrying the mounted slices, they were stained on the slide in a glass container. They were rinsed with double-distilled water twice and then incubated in staining solution for 10 minutes. Afterwards, they were sequentially dehydrated for 4 min/ solution by adding ethanol in ascending concentrations starting with 50% ethanol, 70% ethanol and 95% ethanol. Dehydration with 100% ethanol was performed 4 times for 4 min each. Sections were then cleared with xylene three times for 4 min. Sections could then be coverslipped.

### Gelatin coating of slides

#### TEM

Rats were deeply anesthetised and sacrificed by transaortic perfusion with isotonic saline followed by a mixture of fixative containing 4% paraformaldehyde and 1% glutaraldehyde in 0.1 M phosphate buffer (PB), pH 7.3. Brains were removed and postfixed overnight in the same fixative. Frontal sections (100 µm) cut on a vibratome (Leica VT 1000S) were rinsed in several changes of PB and sections containing septal areas were osmicated in 1% osmium tetroxide in 0.1 M cacodylate buffer for 20 minutes. These sections were further dehydrated using ascending ethyl alcohol concentration steps, followed by two rinses in propylene oxide. Infiltration of the embedding medium was performed by immersing the samples in a 1:1 mixture of propylene oxide and Epon and finally in neat Epon and polymerised at 60°C. Semithin sections (0.5 µm) were prepared for light microscopy, mounted on glass slides, and stained with 1% Toluidine blue. Areas of interest were closely trimmed. Images were acquired with a JEM- 2100Plus Transmission Electron Microscope at 200 kV (Jeol) equipped with a XAROSA CMOS camera (EMSIS).

### Microscopy - Imaging

#### Spinning disk confocal imaging

Fixed confocal imaging was performed using a custom-built inverted microscope equipped with an Olympus IX83 body. For widefield, an LED lamp was used. Excitation lasers used were 405 nm, 488 nm, 561 nm and 640 nm. Lasers were coupled to a CSU-W1 spinning disk unit. Corresponding filters were used and two cameras (sCMOS pco.edge) were used for acquisition. Fixed imaging was performed using a 60x (Olympus, UPLXAPO60XO, 1.42 NA) and a 100x objective.

#### Widefield imaging for Golgi stainings

Golgi images were obtained from a Leica THUNDER Live Cell Imager system equipped with a Leica LED8 Light Engine. Widefield illumination was used for excitation and a Leica DFC 9000GT sCMOS monochrome fluorescence camera. A Leica HC PL APO 20X/0.80 air objective was used.

## 3. Results

### Somatospiny neurons could be identified in the lateral septum through Golgi staining

We first aimed to visualise the somatic spines first described in septal GABAergic neurons (Onteniente et al. 1987), by reproducing initial findings (Robert L. Jakab and Leranth 1990b; 1990a) through comparable methodology. To screen large areas of the septum, without the need for genetic manipulation allowing the observation of an *in vivo* setting, we utilised Golgi-Cox stainings. Here, we could reveal the morphology of somatospiny septal neurons and confirmed their widespread distribution throughout the lateral septum adult rats (Figure 1A). Additionally, we could show that many septal neurons have spiny dendrites, a known feature of somatospiny neurons (**Figure1**A-B).

**Figure 1:**
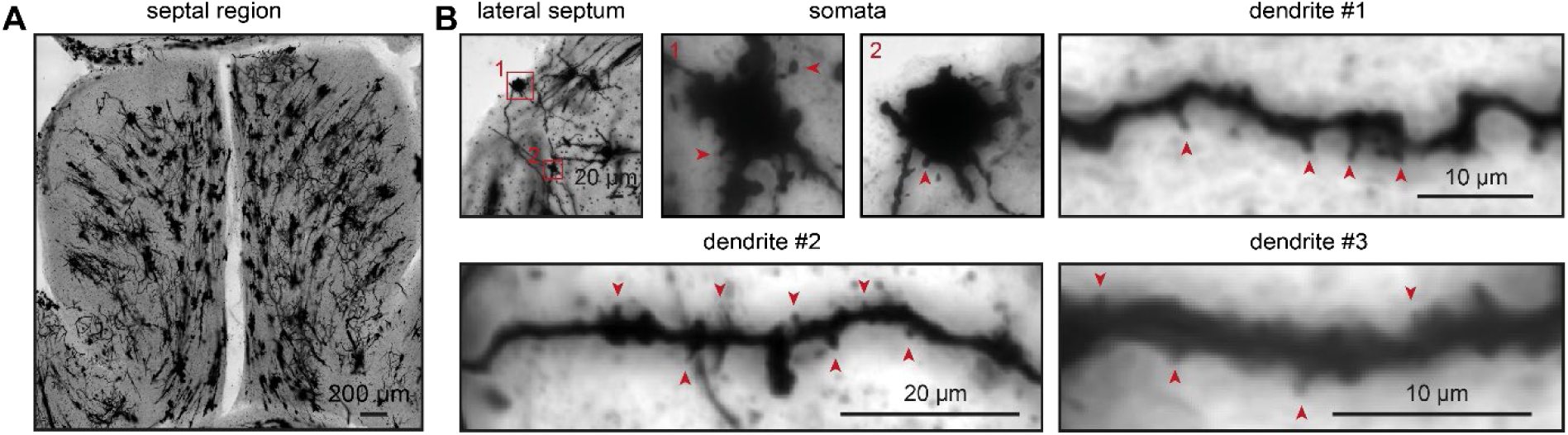
Somatospiny neurons in the rat lateral septum identified by Golgi staining A: Lateral septum of a slice from a Golgi-stained rat brain. Scale bar, 200 µm. B: Representative images of somatospiny neurons found in the lateral septum, two examples of their somata and three of their dendrites. Arrows indicate spines. Scale bars indicated in the images.

### Somatic spines are frequently stubby and smaller than dendritic spines

While Golgi stainings allow for an unbiased approach to observe the morphology of multiple cell types, they come short in the characterisation of molecular features of subcellular structures. To address the question how the ultrastructure of somatic spines was organised, we moved on to high-resolution transmission electron microscopy (TEM). To conclusively confirm that the spine-like structures emerging from neuronal somata are equipped with a postsynaptic density (PSD) and are not merely immature filopodia, we used TEM that does not require a specific postsynaptic marker to label PSDs, allowing an unbiased screening. We could indeed identify somatic spines in the lateral septum of adult rats, thus confirming the presence of somatic spines similar to (Robert L. Jakab and Leranth 1990b) (Figure 2A). Generally, spine synapses are found to be attached to the neuronal dendrite, their attachment to the neuronal soma could have implications on their influence on the cellular output as well as on their composition.

**Figure 2:**
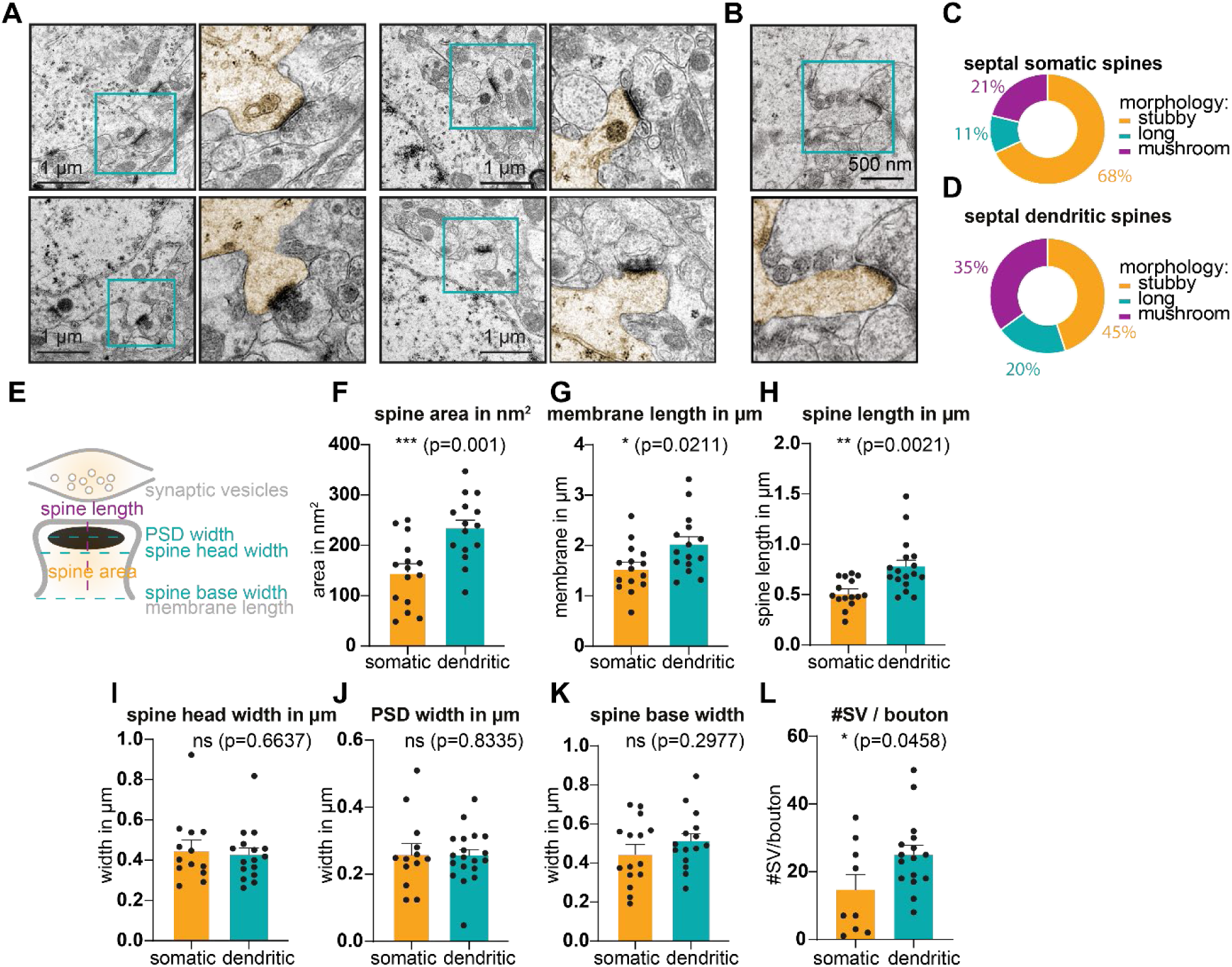
Somatic spines are frequently stubby and smaller than dendritic spines. A, B: Representative transmission electron microscopy images of septal somatic (A) and a dendritic (B) spine of adult rats (N=2) C, D: Spine morphology characterisation of images as shown in (A) and (B), respectively. Morphology grouped into stubby, long filamentous and mushroom spines. E: Schematic depicting spine characterisation measurements depicted in (F)-(L). F-L: Quantification of spine area (F), membrane outlining spines (G), spine length (H), spine head width (I), PSD width (J), width at the base of spines (K) and number of SV on the opposing presynaptic site (L) compared between somatic and dendritic spines. Normal distribution according to Kolmogorov-Smirnov test, unpaired t test. *** p = 0.001 (F), * p = 0.0211 (G), ** p = 0.0021 (H), ns p = 0.6637 (I), ns p = 0.8335 (J), ns p = 0.2977 (K), * p = 0.0458 (L)

To see whether the subcellular localisation of the spines also equipped them with properties that set them apart from their more common dendritic analogues, we compared our findings with characteristics of dendritic spines of the lateral septum (Figure 2B). Since spine morphology can point to the developmental stage or to what kind of input it receives, we first classified and compared the spine morphology of septal somatic and dendritic spines. We observed that most septal spine synapses were stubby spines that have no discernible spine neck, but their spine head rather connects directly to the lumen (Figure 2C-D). 68% of somatic spines were found to be stubby and only 45% of dendritic spines. A large proportion of somatic spines (21%) and of dendritic spines (35%) could be classified as mushroom spines that have a wide spine head on top of a much thinner spine neck connecting them to the lumen. This spine morphology is generally considered to be found in mature spine synapses (Pchitskaya and Bezprozvanny 2020). Finally, 11% of somatic and 20% of dendritic spines were considered long and filamentous that have a prominent spine neck but a thinner spine head than mushroom spines.

We next wondered whether the morphological differences were accompanied by a difference in the volume of somatic spines exceeding that of dendritic spines, measuring multiple characteristics (Figure 2I). Contrary to our expectation, the somatic spines were smaller than their dendritic counterparts in the septum (Figure 2F-G). Accordingly, they were on average 0.5 µm long, with dendritic spines around 0.75 µm (Figure 2H). Despite the clear differences in measured spine area and membrane length, the length of the PSD, spine head and spine neck width at the base of the spine did not differ between somatic and dendritic spines (Figure 2I-K). When comparing the number of synaptic vesicles (SVs) per presynaptic bouton opposed to the postsynaptic density of a spine, we found fewer SVs in boutons opposing somatic than dendritic spines with around 18 SV/bouton for somatic spine synapses and 23 for dendritic spine synapses (Figure 2L).

### Septal spines are mainly excitatory and frequently contain organelles

While dendritic spines are almost exclusively excitatory (McKinney 2010), they can occasionally contain inhibitory PSDs, occasionally at the spine neck (Brusco et al., 2014). We next characterised the spines as excitatory or inhibitory depending on the symmetry of the PSD and the opposing presynaptic site. A similarly electron dense presynapse and PSD were classified as an inhibitory synapse and a clearly more electron dense PSD as excitatory. Synaptic vesicles were found at the presynaptic sites of all classified PSDs, excluding the possibility of having misclassified gap junctions as inhibitory synapses. We found that a vast majority (90%) of somatic spines and all dendritic spines were excitatory in nature (Figure 3A) despite the predominance of inhibitory neurons within the septum.

**Figure 3:**
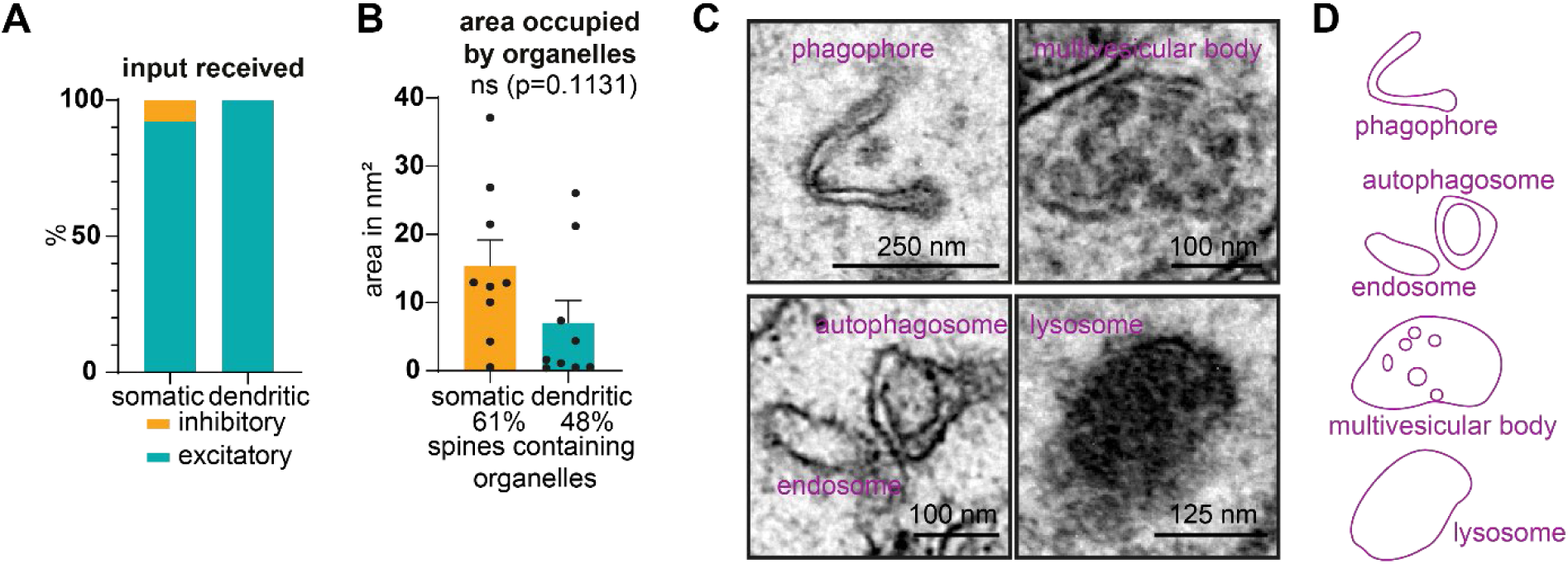
Septal spines are mainly excitatory and frequently contain organelles. A: Share of spines receiving inhibitory or excitatory input as evidenced by a similarly electron dense PSD and presynaptic side for inhibitory synapses and an asymmetrically stronger signal in the PSD for excitatory synapses. B: Percentage of somatic or dendritic spines containing organelles and area occupied by organelles in spines containing organelles. No normal distribution according to Kolmogorov-Smirnov test, Mann Whitney U test. p = 0.1131 C: Additional example organelles found in somatic spines. D: Drawings of organelles seen in (C).

Spine synapse plasticity depends on the availability of various membranous organelles that are generally considered to be shared between dendritic spines that are found in close proximity to each other (Cooney et al. 2002). It is unclear, how the proximity of somatic spines to the organelle dense soma would affect the transport and subsequently the availability of organelles which could affect their plasticity.

We found 60% of all somatic spines contained organelles such as phagophores, multivesicular bodies, endosomes, autophagosomes and lysosomes Figure 3B-D. This suggested that there was a high membrane turnover at somatic septal spines. We did not find any example of a somatic spine containing a spine apparatus, an ER-derived organelle typically found in stable spine synapses (Segal, Vlachos, and Korkotian 2010). Unexpectedly, 50% of dendritic spines were also frequently found to contain various organelles (Figure 3B). A list of organelles found in somatic and dendritic spines synapses is provided (Table 1).

**Table 1:** Percentage of somatic or dendritic spines positive for selected organelles Note that the sum of organelles found can exceed 100% as some spines contained more than one organelle.

|  | somatic spines | dendritic spines |
| --- | --- | --- |
| MVB | 36% | 0% |
| endosome or lysosome | 45% | 0% |
| autophagosome | 18% | 0% |
| ribosomes | 9% | 55% |
| spine apparatus | 0% | 18% |
| other | 9% | 36% |

#### Development of somatic spines occurs independently of extra-septal input in dissociated septal cultures

Although it had been suggested that septal somatic spines receive hippocampal input (Robert L. Jakab and Leranth 1990a), it is presently unclear, whether it is required for the somatic spines to develop or whether the somatospiny neurons are able to form somatic spines in the absence of hippocampal input. To investigate this question, we next set out to establish methods to be used to visualise somatic spines in the lateral septum that would allow us to easily manipulate them and characterise their content. One optically easily accessible model system in which this would be feasible, are dissociated primary cultures that are commonly used, especially for hippocampal and cortical cells (Meberg and Miller 2003; Sahu et al. 2019). This culturing system (Figure 4A-B) allowed us to effectively image the outlines of complete neuronal somata in combination with a multitude of synaptic markers. Here, we could combine genetic manipulation with subsequent immunostainings to understand the molecular composition of these somatic spines during development.

**Figure 4:**
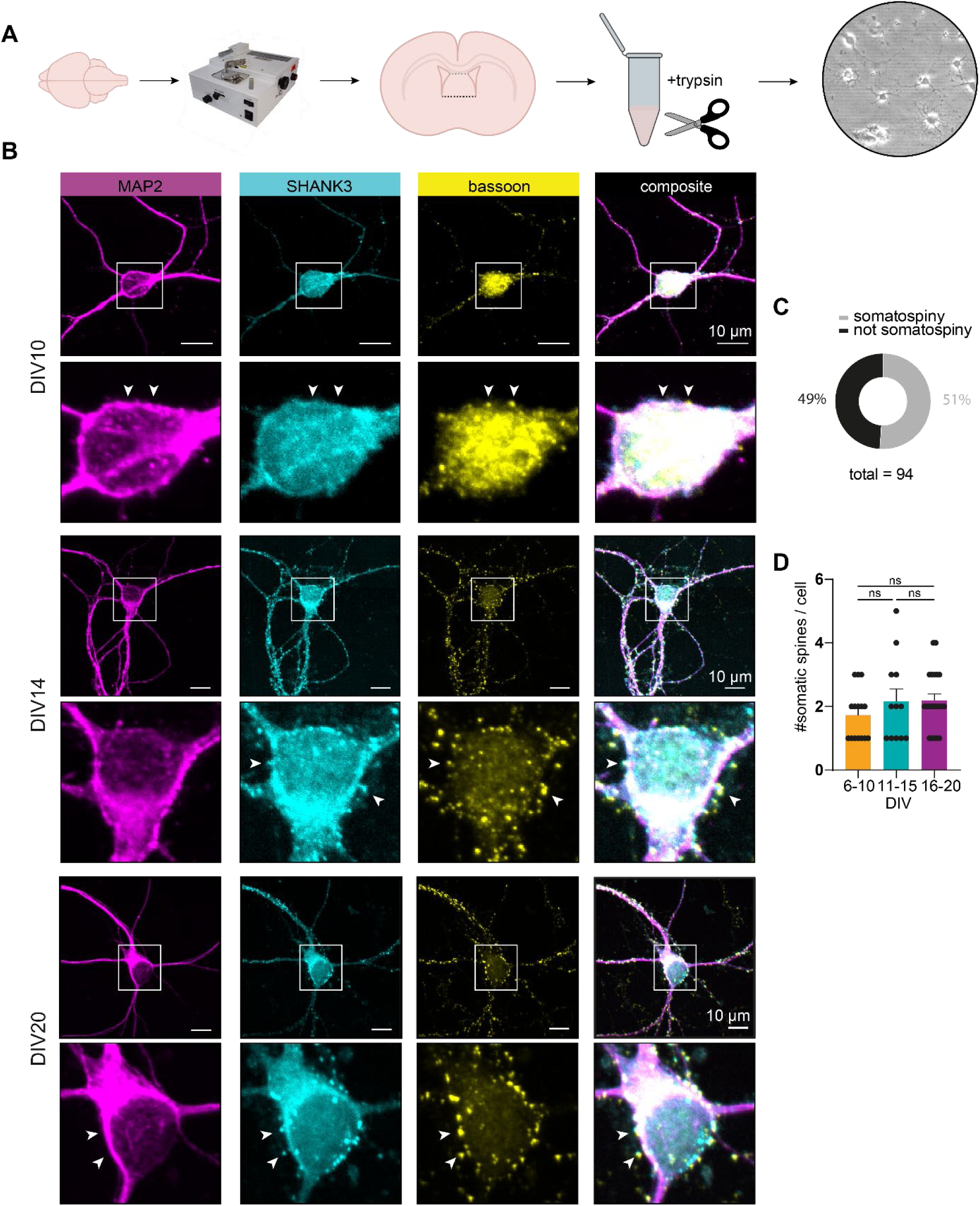
Somatospiny neurons develop independently of specific input in dissociated primary cultures A: Schematic showing dissection method used to prepare septal dissociated cultures B: Dissociated septal neurons fixed at selected DIV, immunostained against neuronal, somatodendritic marker MAP2 (magenta), postsynaptic scaffolding protein SHANK3 (cyan) and presynaptic marker bassoon (yellow). Scale bar, 10 µm. Indicated zoom-ins of somata below the image. Somatic spines indicated by arrows. C: Percentage of neurons in septal dissociated culture stained as in B that bear somatospines (somatospiny) or not. n = 94 cells (N=11) D: Number of somatic spines at different stages in development found on somatospiny neurons as in B. Mann Whitney U test: ns DIV6-10 vs DIV11-15 p = 0.5360, ns DIV11- 15 vs DIV16-20 p = 0.6822

On average, we found that 51% (n=48 of 94 cells) of septal neurons were somatospiny. This conclusion was supported by the presence of synaptic markers outside the bounds of the somatodendritic marker MAP2-signal (Figure 4C). On average we observed two somatic spines on somatospiny neurons with no significant differences between days *in vitro* (DIV)6 and 20 (Figure 4D).

This confirmed the ability of somatospiny neurons to develop somatic spines in the absence of extra-septal e.g. hippocampal input.

#### Somatic spines observed in organotypic septal slices show classical postsynaptic markers

While dissociated cultures are especially accessible, specific connectivity patterns between cells are lost. To keep a similar level of accessibility, but simultaneously increase the level of complexity and move closer to the native connectivity, we next aimed to generate organotypic septal slices from postnatal rats (postnatal day (P) 5-7) (Figure 5A). To identify a number of neurons, we sparsely labelled neurons using field electroporation of plasmid DNA encoding mRuby2 used as a volume marker (Figure 5B). We then quantified the number of somatic spines found on a somatospiny neurons as well as the share of somatospiny neurons and compared our findings for different developmental stages (DIV10-29). We found that 26% (n=48 of 187 cells) of all electroporated neurons were somatospiny (Figure 5C). In contrast to dissociated cultures, we found an increase in the number of somatic spines on a somatospiny neuron during development (Figure 5D). At DIV10, only around 2 somatic spines could be found on a somatospiny neuron, increasing to ∼6 at DIV17-22 and to ∼9 at DIV29 (Figure 5D).

**Figure 5:**
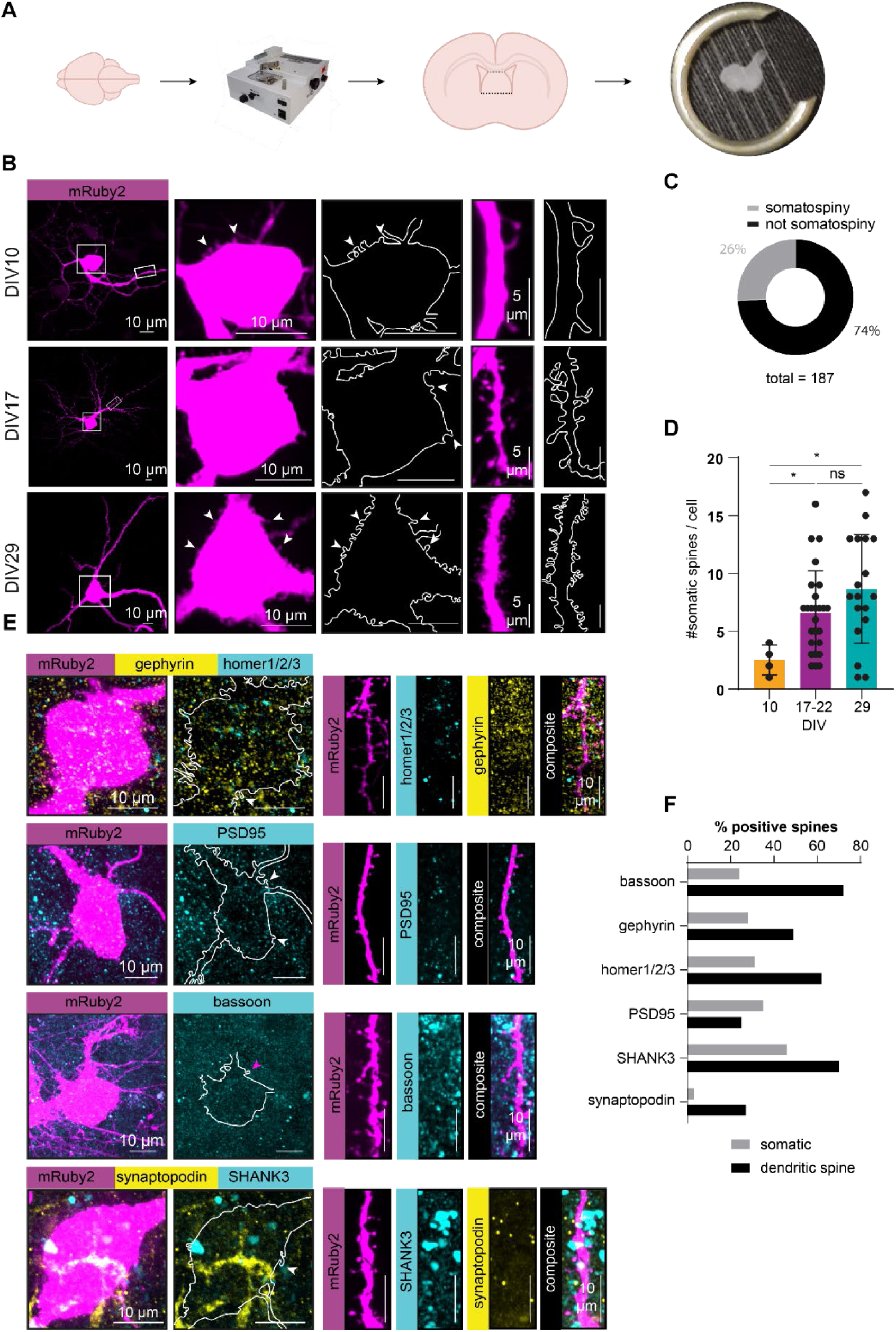
Somatic spines contain classical postsynaptic markers in organotypic septal slice cultures A: Schematic showing dissection method used to prepare septal organotypic cultures B: Maximum-projections of electroporated septal neurons expressing the cell fill mRuby2 fixed at selected DIV. Scale bar 10 µm for overview and somatic images, 5 µm for dendrites. Outlines drawn in panels right next to cell fill images. C: Percentage of somatospiny neurons in septal organotypic culture and those without somatic spines. n = 187 cells (N=15) D: Number of somatic spines at different stages in development found on somatospiny neurons as in (B). Mann-Whitney U test: * DIV10 vs. DIV17-22 p = 0.0142, ns DIV17- 22 vs. DIV29 p = 0.1039, * DIV10 vs DIV29 = 0.0187 E: Electroporated septal slices fixed at DIV17-29, immunostained for selected synaptic markers. F: Quantification of somatic spines containing selected synaptic markers as in (E).

After confirming the validity of organotypic septal slices for the characterisation of somatic spines, we sought to identify a common synaptic marker for the somatic spines. We thus stained for common synaptic markers, including the universal, presynaptic scaffolding protein bassoon, the inhibitory postsynaptic scaffolding protein gephyrin and multiple excitatory postsynaptic proteins: homer1/2/3, PSD95 and SHANK3. Despite most somatic spines displaying mushroom or stubby spine-like morphology, we found that many of them do not contain classical synaptic markers (Figure 4F). We found around 24% of spines that were positive for the presynaptic scaffolding protein bassoon (Figure 4F). Around 28% of spines were found to contain the inhibitory scaffolding protein gephyrin, 31% homer1/2/3 and 35% PSD95 (Figure 4F). Around 46% of somatic spines were positive for excitatory scaffolding protein SHANK3. To validate our synaptic markers, we also quantified the abundance of septal dendritic spines positive for the same markers (Figure 4E). Where we found that 80% of spines were positive for the universal presynaptic marker bassoon. 49% contained gephyrin, 62% homer 1/2/3 and only 25% PSD95. 70% were positive for SHANK3

## 4. Discussion

Here, we reaffirm the existence of septal somatic spines in the lateral septum. New approaches allowed us to identify the molecular composition of septal somatic spines and to compare them to dendritic spines of the lateral septum.

We first set out to classify whether all septal somatic spines were excitatory in nature. While dendritic spines are generally excitatory and known as the major excitatory input processor of the neurons (Yuste and Bonhoeffer 2004), somatic spines are less well characterised. With the help of transmission electron microscopy (TEM), we found that virtually all septal somatic spine synapses are excitatory while all septal dendritic spines are excitatory. To identify molecular markers for somatic spines, we used dissociated cultures and organotypic slices, both of which are disconnected from their native network. While this reduced preparation results in a network that lacks input from other brain regions, such methods gave us the opportunity to study whether septal somatic spine formation existed as a postsynaptically built-in programme of somatospiny septal neurons. Our *in vitro* work showed that somatic spines could develop without requiring input from any other brain region than the septum. This indicated that the somatic spines developed independently of excitatory input e.g from the hippocampus.

*In vivo* somatic spines receive input from hippocampal somatostatin neurons or catecholaminergic input as suggested previously (Robert L. Jakab and Leranth 1990a; R L Jakab and Leranth 1991). This input could be replaced depending on available cells in *in vitro* systems as in (Agi et al. 2024) where it has been shown that axons self-sort depending on available postsynaptic targets. To answer the question whether *in vivo* septal somatic spines receive synaptic input from the hippocampus, tracing experiments would be most suitable.

Additionally, the high density of synaptic markers in septal organotypic slices indicated a mature and healthy environment, but made it challenging to judge whether an individual somatic spine was positive for a specific marker. Analysis of dendritic spines in the same slices however proved that the vast majority was positive for the universal presynaptic scaffolding protein bassoon and often other established synaptic markers. This indicated that the low share of somatic spines positive for the selected markers was not caused by methodological shortcomings, but reflected a decreased content of established synaptic proteins, possibly indicating that spine synapses were less mature than those observed with TEM. While we tested a number of different synaptic markers, we were not able to use more than two at the same time. Thus, our results could also indicate a high variability in the molecular composition of somatic spines. A more complete overview could be achieved by patch-sequencing and thanks to recent advances - additional single-cell proteomics - but would be challenging to achieve due to the lack of a somatospiny neuron-specific marker and the predominance of somatodendritic and axonal proteins in the resulting sample.

We were particularly interested in the occupancy of dendritic and somatic spines by different sets of organelles. Here, using electron microscopy, we are able to identify the precise organelle through the detection of their outline. While only depicting a snapshot in the life of a spine synapse, this method does not rely on the various marker proteins that are rarely exclusive to a specific type of organelle but rather show gradients across their development. It thus allowed us to confidently identify multiple organelles involved in the turnover of membranes and the degradation of proteins. The difference in organelles present pointed to a different underlying physiology and membrane turnover, associated with increased memory capacity (Frank et al. 2018). On a subcellular level, it also indicated that organelle transport was organised differently, despite the functional similarity - as indicated by the similar appearance of the PSD - of the target structures. To our surprise, we found that septal somatic spines were smaller than septal dendritic spines. This was in contrast to somatic spines containing organelles more often while individual area occupied by organelles was comparable between the different spine types.

Our TEM quantification was based on electron density of the postsynaptic density that exhibits a stronger signal at excitatory postsynapses, causing the synapse to appear asymmetric when comparing the pre- and postsynaptic site. To gain an additional level of certainty, immunogold labelling could be performed using antibodies raised against proteins typically found at the excitatory postsynapse such as PSD95 and homer or those typically found at the inhibitory postsynapse such as gephyrin.

Even prior to TEM, Golgi stainings are arguably the oldest method of visualising neurons in the brain (Golgi 1886) and can still serve as a method to do so reliably. This method is, however, prone to artefacts that are often visible as black dots of ∼1 µm diameter, similar to that of spine synapses. This already caused Camillo Golgi to hypothesise that dendritic spines are staining artefacts and not structures found on dendrites (Golgi 1886). While it is undisputed that dendritic spine synapses exist and are quite clearly visible on dendrites of Golgi-stained samples, somatic spines are less easily distinguished from artefacts surrounding the neuronal somata. We could find evidence of somatic spines of adult rats in Golgi stainings, we did however not quantify their occurrence since not every assumed somatic spine was undoubtedly a spine rather than a staining artefact. In combination with other methods, we can however confidently say that the adult rat lateral septum contains a vast number of somatospiny neurons. For further investigation, we decided to focus on fluorescent microscopy and TEM to visualise single neurons and the composition of the somatic spines in multi- colour images.

## Conclusion

We confirm the existence of somatic spines in the lateral septum and characterise them in different model systems where they can develop independently from input from other brain regions than the septal area. They are smaller than dendritic spines but more frequently contain membranous organelles. Despite being excitatory in nature, they are generally negative for most excitatory synaptic markers, but show a similar developmental timeframe to dendritic spines. While the investigation of somatic spines has subsided in the last few years, the lateral septum is not the only region containing somatospiny neurons. Notably, they are found on ganglionic neurons in the peripheral nervous system in the hypothalamus, on Purkinje cells in the cerebellum and in adult hippocampus (Hirano, Dembitzer, and Yoon 1977; Ifft and McCarthy 1974; Shoop et al. 1999; Wenzel et al. 1994). The new approaches established here and molecular characterisations of somatic spines, provide a comprehensive foundation to further elucidate the enigmatic somatic spines of septal neurons.

## Acknowledgements

This work was supported by the Deutsche Forschungsgemeinschaft (DFG) SFB1315 A03 and the Excellence Strategy – EXC- 2049–390688087 to MB and MM.

